# Novel neurofilament light (*Nefl*) E397K mouse models of Charcot-Marie-Tooth type 2E (CMT2E) present early and chronic axonal neuropathy

**DOI:** 10.1101/2025.02.02.636117

**Authors:** Dennis O. Pérez-López, Audrey A. Shively, F. Javier Llorente Torres, Mohammed T. Abu-Salah, Michael L. Garcia, W. David Arnold, Monique A. Lorson, Christian L. Lorson

## Abstract

Charcot-Marie-Tooth (CMT) is the most common hereditary peripheral neuropathy with an incidence of 1:2,500. CMT2 clinical symptoms include distal muscle weakness and atrophy, sensory loss, toe and foot deformities, with some patients presenting with reduced nerve conduction velocity. Mutations in the neurofilament light chain (*NEFL*) gene result in a specific form of CMT2 disease, CMT2E. *NEFL* encodes the protein, NF-L, one of the core intermediate filament proteins that contribute to the maintenance and stability of the axonal cytoskeleton. To better understand the underlying biology of CMT2E disease and advance the development of therapeutics, we generated a *Nefl^+/E397K^* mouse model. While the *Nefl^+/E397K^* mutation is inherited in a dominant manner, we also characterized *Nefl^E397K/E397K^* mice to determine whether disease onset, progression or severity would be impacted. Consistent with CMT2E, lifespan was not altered in these novel mouse models. A longitudinal electrophysiology study demonstrated significant in vivo functional abnormalities as early as P21 in distal latency, compound muscle action potential (CMAP) amplitude and negative area. A significant reduction in the sciatic nerve axon area, diameter, and G-ratio was also present as early as P21. Evidence of axon sprouting was observed with disease progression. Through the twelve months measured, disease became more evident in all assessments. Collectively, these results demonstrate an early and robust in vivo electrophysiological phenotype and axonal pathology, making *Nefl^+/E397K^* and *Nefl^E397K/E397K^*mice ideal for the evaluation of therapeutic approaches.

## Introduction

Charcot-Marie-Tooth (CMT) is one of the most common inherited neurological disorders with an incidence of ∼1:2,500. Clinical classification of CMT is primarily based on patient electrophysiology measurements. There are eight classifications of CMT (1-7 and CMT-X). CMT1 and CMT2 are the most prevalent forms and are associated with deficiencies in myelination or axonal dysfunction and degeneration, respectively. CMT2 is a result of mutations in many genes. CMT2 clinical symptoms are characterized as slow but progressive with a wide spectrum of distal muscle weakness and atrophy, sensory loss, decreased deep-tendon reflexes, toe, and foot deformities, gait disturbances, with some patients presenting with reduced motor nerve conduction velocity (MNCV). CMT2E is caused by mutations in the neurofilament light chain (*NEFL*) gene and is primarily inherited in an autosomal dominant manner. CMT2E disease onset and severity are variable even within families with the same mutation (1–6).

Neurofilaments, also known as intermediate filaments, contribute to the expansion and maintenance of axonal caliber, axon structure and transport. Mutations in neurofilaments lead to axonal dysfunction, characteristic of CMT2. In peripheral nerves, neurofilament light protein (NF-L) neurofilament heavy (NF-H), neurofilament medium (NF-M) and peripherin assembly to form intermediate filaments. In the central nervous system (CNS), intermediate filaments are composed of the core proteins NF-H, NF-M, NF-L, and alpha internexin (3, 7–11). There are over 30 different mutations in *NEFL* that are distributed throughout the three functional domains (head, rod, tail). Homozygous recessive mutations in *NEFL*, while rare, have been reported; however, CMT2E is primarily associated with dominant mutations. The most prevalent *NEFL* mutation, *E396K*, is located within a highly conserved motif at carboxy terminal end of the rod domain. The rod domain is involved in dimerization and mutations, such as *E396K*, likely disrupt neurofilament assembly (12). Mutations in *NEFL* result in impaired axonal assembly, accumulation of neurofilaments, atrophied axons, and perturbed localization and transport of the mitochondria (13–15).

There are several mouse models of *Nefl* that are associated with patient mutations (*Nefl^N98S^*, *Nefl^P8R^*, *Nefl^L394P^*, h*NEFL^E396K^* and h*NEFL^P22S^*) (14, 16–21). *Nefl^N98S^* and *Nefl^P8R^*models have the orthologous mutation generated within the mouse genome while the *Nefl^L394P^*mouse has mutant mouse *Nefl* driven by the murine sarcoma virus promoter. h*NEFL^E396K^* and h*NEFL*^P22S^ models each overexpress the human *E396K* or *P22S* transgenes, respectively, with wild type mouse *Nefl* expression present. Each model recapitulates many phenotypes associated with CMT2E disease to varying degrees. The *Nefl^N98S^* mouse presents with the earliest disease onset and most robust axon pathology (∼6 weeks) (16, 22). As *NEFL^E396K^* is a predominant mutation, we wanted to examine disease biology in the context of the mouse genome where the orthologous mutation (*E397K*) is present. While *NEFL* mutations are predominantly dominant, we examined *Nefl^+/E397K^* and *Nefl^E397K/397K^* mice to determine whether disease onset or severity were altered. Here we report our electrophysiological findings, axonal pathology and neuromuscular junction innervation status for mouse models, *Nefl^+/E397K^* and *Nefl^E397K/397K^*. These models show a clinically relevant phenotype with a chronic axonopathy characterized by electrophysiological deficits. Importantly, these mouse models provide an important context for future therapeutic development.

## Results

### Generation of the *Nefl* E397K mouse models of CMT2E

Mutations in the *NEFL* gene result in CMT2E, a slow progressive disease characterized by motor function deficits and axonal pathology. To better understand disease development and mechanistic pathways, we generated a *Nefl^+/E397K^* mouse model with an orthologous mutation that corresponds to the human *NEFL-E396K* mutation (**Fig. 1A**). While *Nefl-E397K* presents as a dominant mutation, we wanted to determine whether the *Nefl^E397K/E397K^* mouse would be more severe; therefore, *Nefl^+/E397K^* and *Nefl^E397K/E397K^*mice were analyzed. Mice were genotyped using quantitative PCR. Survival was monitored over 360 days with 100% survival for the *Nefl^+/E397K^* and the *Nefl^E397K/E397K^* mice (**Fig. 1B**), consistent with CMT2E patient outcomes. Weight was measured from P1-P360; there was statistical difference between wild type (mean 27.74 grams) and *Nefl^+/E397K^* mice (mean 28.72 grams, *P*<0.0001) as well as wild type and *Nefl^E397K/E397K^* mice (mean 25.69 grams, *P*<0.0001). From ∼P240, *Nefl^E397K/E397K^* mice trended towards weighing less (**Fig. 1C).** There were no sex-based differences for weight; all male mice gained weight at higher rates than females of the same genotype (not shown). *Nefl^+/E397K^* and the *Nefl^E397K/E397K^*mice were phenotypically indistinguishable compared to their wild type littermates at early time points; however, as disease progressed there were overt signs of hindlimb weakness (**Fig. 1D** and not shown).

**Figure 1.**
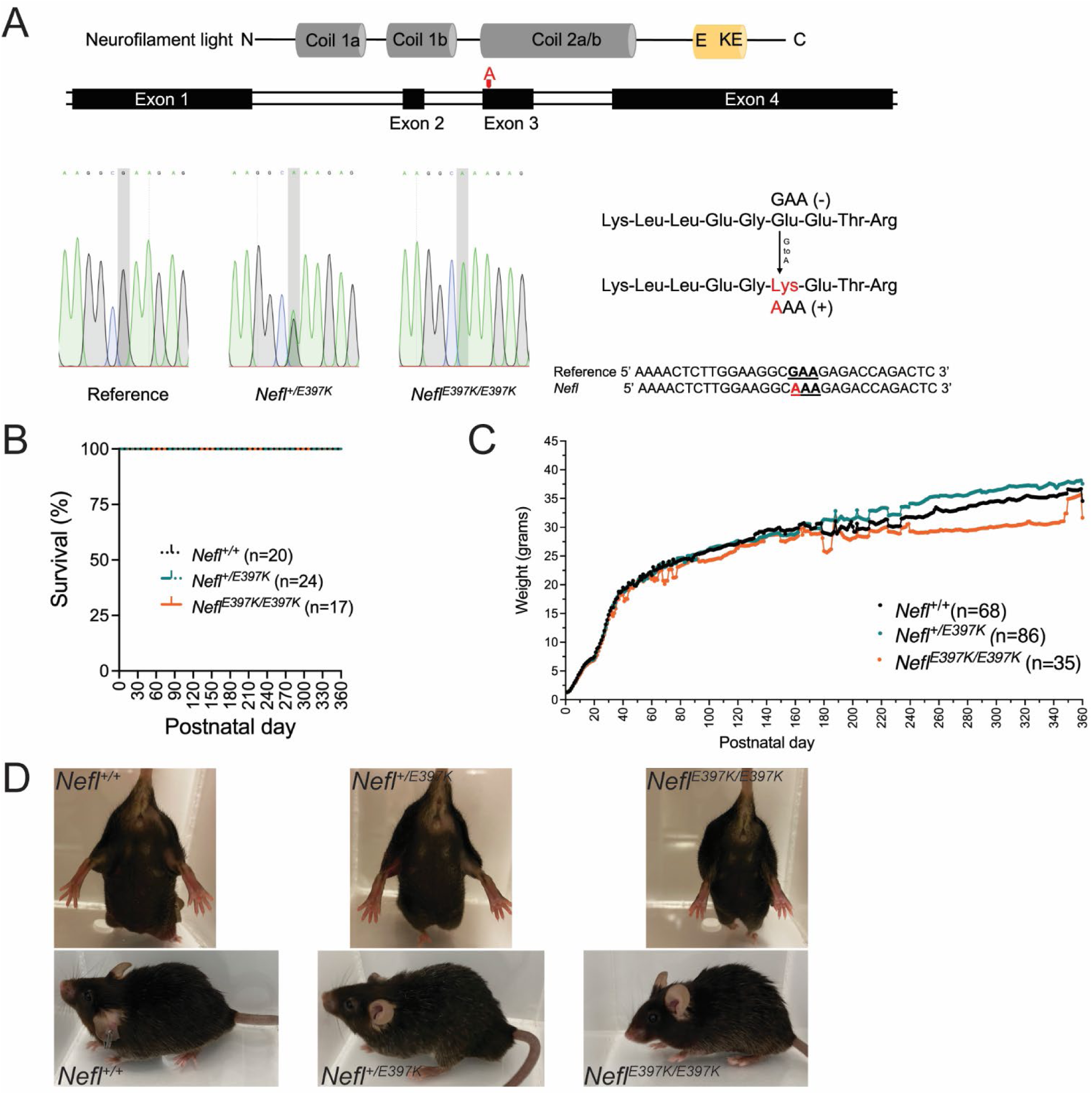
Generation of the *Nefl* E397K mouse models of CMT2E. (A) Cartoon depicting the neurofilament light protein (NF-L). The glutamic acid to lysine (E397K) alteration with nucleotide changes GAA to AAA were generated using CRISPR Cas9. The wild type *Nefl*, *Nefl*^+/E397K^ and *Nefl^E^*^397K/E397K^ sequences generated from tail DNA isolated from wild type and *Nefl* mutant mice. (B) Percent survival for wild type (black), *Nefl*^+/E397K^ (teal) and *Nefl*^E397K/E397K^ (orange) mice recorded from P0 to P360. (C) Weight in grams for wild type (black), *Nefl*^+/E397K^ (teal) and *Nefl^E^*^397K/E397K^ (orange) mice recorded from P0 to P360. One way ANOVA with Tukey’s multiple comparison test was used to determine significance. (D) Images of wild type and *Nefl* mutant mice. Top images are of female mice at P120. Bottom images are of male mice at P360. N=number of animals evaluated, g= grams, P=postnatal day.

### Electrophysiology showed an early clinically relevant phenotype

One of the hallmarks of CMT2E disease is the presence of distinct electrophysiological abnormalities, which can also be assessed in mice. While there is variability within the patient population, CMT2E patients have relatively preserved MNCV values with reduced amplitudes of sensory and CMAP responses. However, a subset of CMT2E patients have reduced MNCV that is attributed to a decrease in axon caliber and not myelination (2, 23, 24).

An initial electrophysiology study was conducted on four cohorts of animals at three weeks, twelve weeks, six months and twelve months of age. At each time point, animals were sacrificed and tissues collected. Electrophysiological measurements quantitatively measured stimulation of the sciatic nerve and response of the gastrocnemius muscle. The in vivo axon caliber (distal latency), CMAP (peak-to-peak (P/P) amplitude) and activated muscle fibers by the stimuli (negative area) were measured. At three weeks, distal latency was significantly prolonged in *Nefl^+/E397K^* (0.7586ms, *P*=0.0026) and *Nefl^E397K/E397K^* (0.7963ms, *P*=0.0085) mice when compared to wild type (0.5271ms) (**Fig. 2A**). CMAP amplitude and negative area were also significantly different in the *Nefl* mutants when compared to the wild type cohort (**Fig. 2A**). At twelve weeks, the distal latency of *Nefl^+/E397K^* (0.7532ms, *P*=0.0008) and *Nefl^E397K/E397K^* (0.9213ms, *P*<0.0001) mice was prolonged compared to wild type mice (0.5208ms) with significant worsening in *Nefl^E397K/E397K^* mice (**Fig. 2B**). CMAP amplitude also remained significantly different between the *Nefl* mutants and the wild type cohort at twelve weeks (**Fig. 2B**). Interestingly, at twelve weeks there was improvement in the negative area of *Nefl^+/E397K^* mice but *Nefl^E397K/E397K^*mice remained significantly different from wild type mice (**Fig. 2B**). At six and twelve months, there remained significant differences between wild type and *Nefl* mutants in distal latency; however, CMAP amplitude and negative area values were similar between all cohorts (**Fig. 2C-D**). These results show that *Nefl^+/E397K^* and *Nefl^E397K/E397K^* mice have significant defects in distal latency starting at P21 and continuing throughout the study (P360).

**Figure 2.**
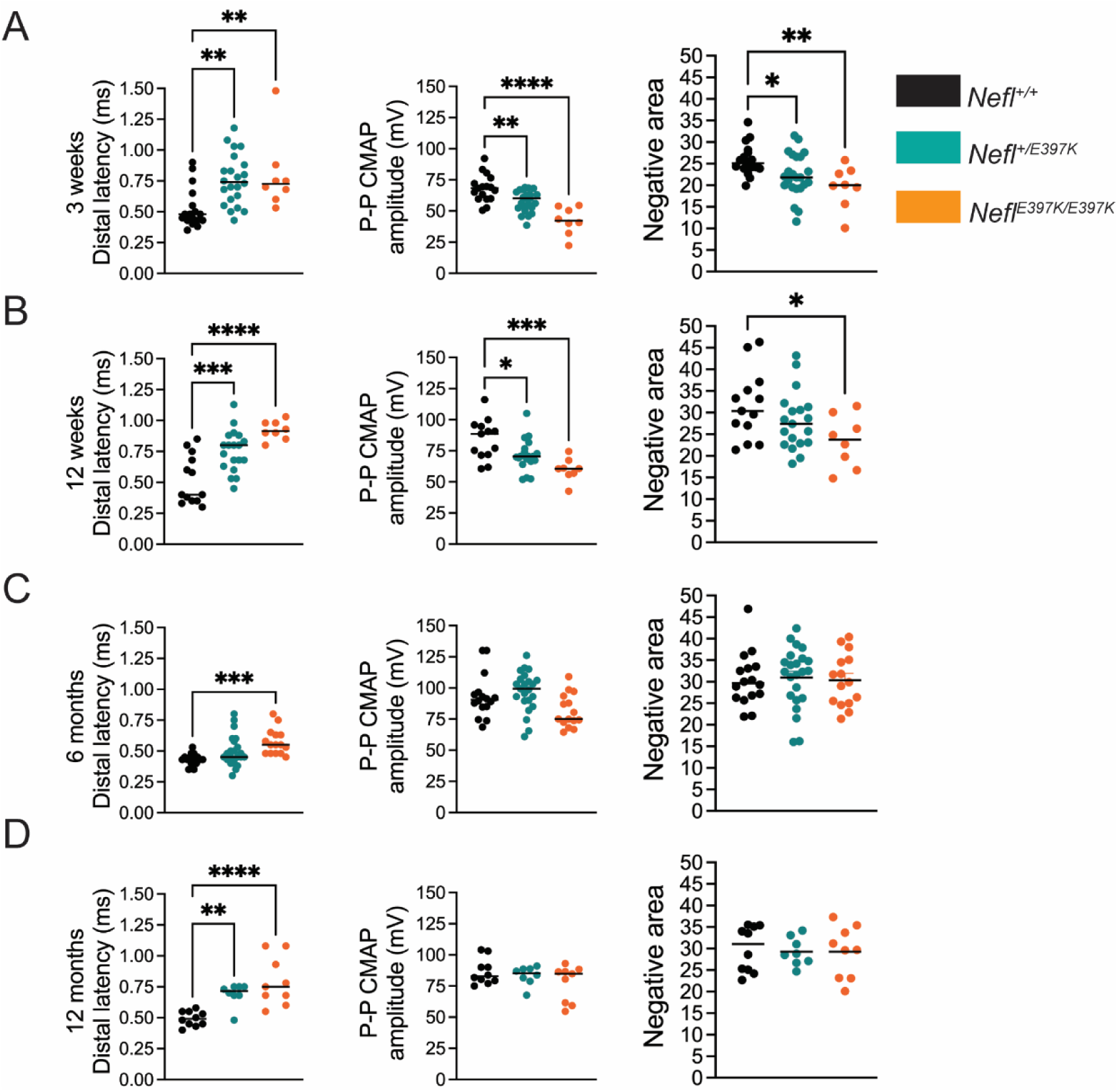
Electrophysiology showed an early clinically relevant phenotype. Distal latency, Peak to Peak (P-P) CMAP amplitude and negative area were measured following stimulation of the sciatic nerve and recordings from the gastrocnemius muscle. Wild type (black), *Nefl*^+/E397K^ (teal) and *Nefl^E^*^397K/E397K^ (orange) mice were evaluated at three weeks, twelve weeks, six months, and twelve months. (A) Electrophysiology recordings at three weeks, wild type (N=17), *Nefl*^+/E397K^ (N=23) and *Nefl^E^*^397K/E397K^ (N=8). Distal latency for wild type (mean=0.5271), *Nefl*^+/E397K^ (mean=0.7586, *P*=0.0026) and *Nefl^E^*^397K/E397K^ (mean=0.7963, *P*=0.0085) mice. P-P CMAP amplitude for wild type (mean=68.13), *Nefl*^+/E397K^ (mean=57.82, *P*=0.0037) and *Nefl^E^*^397K/E397K^ (mean=42.23, *P*<0.0001) mice. Negative area for wild type (mean=25.89), *Nefl*^+/E397K^ (mean=22.30, *P*=0.0341) and *Nefl^E^*^397K/E397K^ (mean=19.65, *P*=0.0051) mice. (B) Electrophysiology recordings at twelve weeks, wild type (N=13), *Nefl*^+/E397K^ (N=19) and *Nefl^E^*^397K/E397K^ (N=8). Distal latency for wild type (mean=0.5208), *Nefl*^+/E397K^ (mean=0.7532, *P*=0.0008) and *Nefl^E^*^397K/E397K^ (mean=0.9213, *P*<0.0001) mice. P-P CMAP amplitude for wild type (mean=84.15), *Nefl*^+/E397K^ (mean=71.43, *P*=0.0224) and *Nefl^E^*^397K/E397K^ (mean=59.93, *P*=0.0005) mice. Negative area for wild type (mean=31.63), *Nefl*^+/E397K^ (mean=28.08, NS) and *Nefl*^E397K/397K^ (mean=23.34, *P*=0.0240) mice. (C) Electrophysiology recordings at six months, wild type (N=16), *Nefl*^+/E397K^ (N=23) and *Nefl^E^*^397K/E397K^ (N=15). Distal latency for wild type (mean=0.4338), *Nefl*^+/E397K^ (mean=0.4983, NS) and *Nefl^E^*^397K/E397K^ (mean=0.5727, *P*=0.0007) mice. P-P CMAP amplitude and negative area were not statistically significant between genotypes. (D) Electrophysiology recordings at twelve months, wild type (N=10), *Nefl*^+/E397K^ (N=8) and *Nefl^E^*^397K/E397K^ (N=9). Distal latency for wild type (mean=0.4920), *Nefl*^+/E397K^ (mean=0.6900, *P*=0.0065) and *Nefl^E^*^397K/E397K^ (mean=0.7922, *P*<0.0001) mice. P-P CMAP amplitude and negative area were not statistically significant between genotypes. Statistical significance was determined used ordinary one-way ANOVA and Dunnett’s multiple comparisons test. Ms=milliseconds, mV=millivolts, CMAP= compound muscle action potential, NS=not significant, N=number of mice evaluated.

The initial electrophysiology study showed prolonged distal latency from three weeks to twelve months in the *Nefl* mutants; however, CMAP amplitude and negative area differences improved from P21 to P360. To determine whether these changes reflected disease progression, a twelve-month longitudinal study was performed on a cohort of wild type, *Nefl^+/E397K^*and *Nefl^E397K/E397K^* mice. The longitudinal assessments showed that the *Nefl^E397K/E397K^* presented with prolonged distal latency as early as P21, while *Nefl^+/E397K^* distal latency became statistically different from wild type mice at P90 (**Fig. 3A**). The mean distal latency between wild type (0.4630ms) and *Nefl^+/E397K^*mice (0.6545ms) was statistically different (*P=*0.0002), as well as between *Nefl^E397K/E397K^* mice (0.8745ms, *P*<0.0001). There was also a statistical significance in distal latency between *Nefl^+/E397K^*and *Nefl^E397K/E397K^* mice (*P=*0.0010). *Nefl^E397K/E397K^*distal latency was statistically different from wild type mice throughout the study (P21-P360) suggesting, like some CMT2E patients, there is a reduction in axon caliber (**Fig. 3A**). Interestingly, P21-P60 *Nefl^E397K/E397K^* CMAP P-P amplitude values were statistically different from wild type mice; however, for the remainder of the study differences varied (**Fig. 3B**). There was statistical difference between the mean CMAP amplitude values for wild type (81.12mV) and *Nefl^+/E397K^* mice (76.41mV, *P*=0.0319) and *Nefl^E397K/E397K^*mice (66.39mV, *P*=0.0001). As well, there were CMAP P-P amplitude differences between *Nefl^+/E397K^* and *Nefl^E397K/E397K^*mice (*P=*0.0012). The same trends were observed for negative area measurements in *Nefl^E397K/E397K^* mice (**Fig. 3C**). There was no statistical difference between the mean negative area values for wild type (26.55) and *Nefl^+/E397K^*mice (26.51) and *Nefl^E397K/E397K^* mice (23.93). Importantly, results of the longitudinal study were consistent with the mixed cohort electrophysiology study; distal latency was severely impacted early in *Nefl^E397K/E397K^*mice with differences occurring later in *Nefl^+/E397K^* mice. In *Nefl^E397K/E397K^* and *Nefl^+/E397K^* mice, CMAP amplitude and negative area differences were apparent early; however, differences were not measured later in the disease. The subtle differences between the two studies were likely attributed to disease variation between the cohorts.

**Figure 3.**
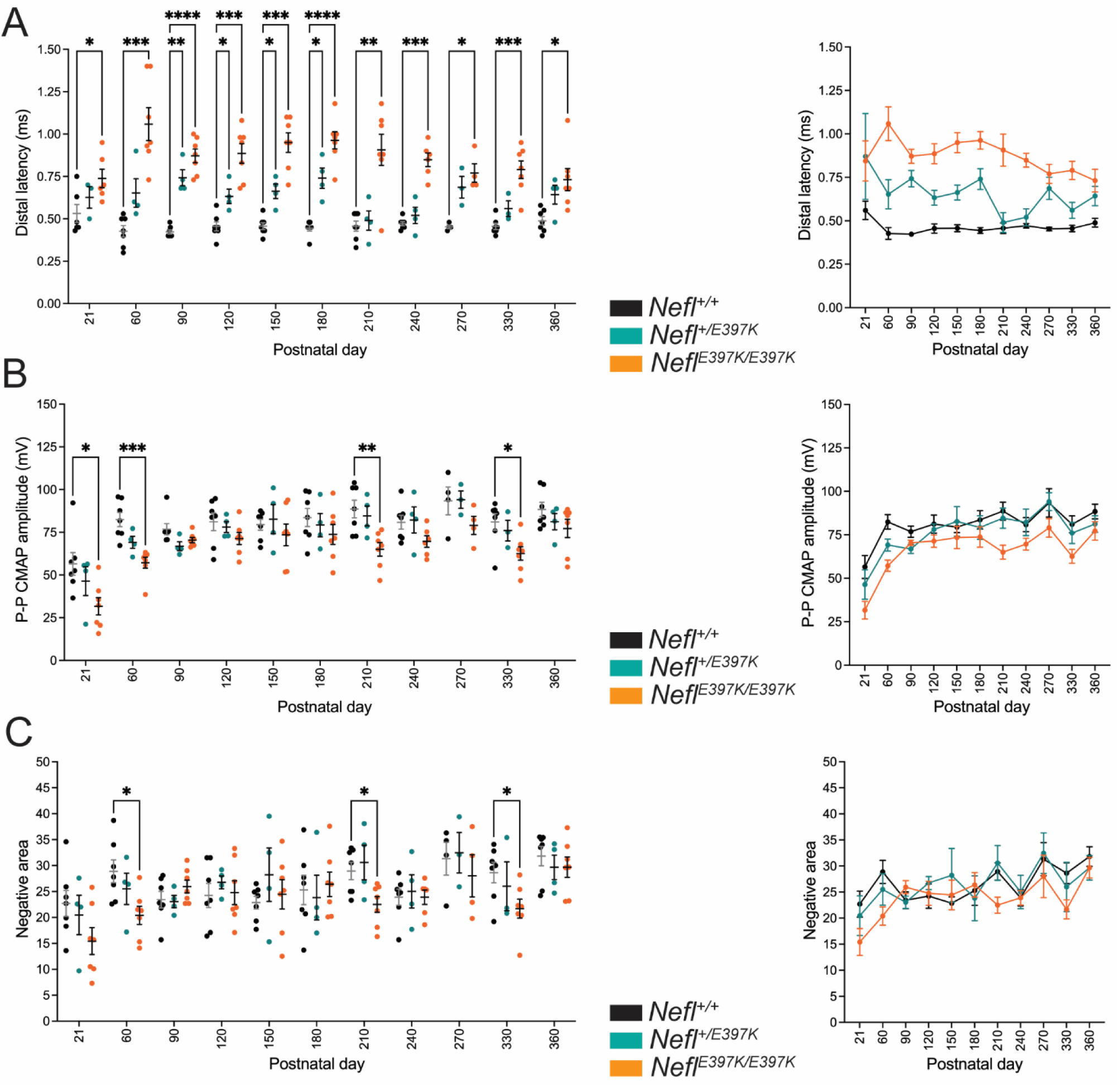
Longitudinal electrophysiology study of wild type and *Nefl* mutant mice. Distal latency, Peak to Peak (P-P) CMAP amplitude and negative area were measured following stimulation of the sciatic nerve and recordings from the gastrocnemius muscle. Wild type (black, N=7), *Nefl*^+/E397K^ (teal, N=4) and *Nefl^E^*^397K/E397K^ (orange, N=7) mice were evaluated at P21, P60, P90, P120, P150, P180, P210, P240, P270, P330 and P360 days. (A) Distal latency *P*\*=0.0148-0.0395, *P*\*\*=0.0040-0.084, *P*\*\*\*=0.0001-0.0007, *P*\*\*\*\*<0.0001. Wild type (mean=0.4630), *Nefl*^+/E397K^ (mean=0.6545, *P*=0.0002), *Nefl^E^*^397K/E397K^ (mean=0.8745, *P*<0.0001), *Nefl*^+/E397K^ mean compared to *Nefl^E^*^397K/E397K^ mean (*P*=0.0010). (B) Peak-Peak CMAP amplitude *P*\*=0.0229-0.0242, *P*\*\*=0.0074, *P*\*\*\*=0.0010. Wild type (mean=81.12), *Nefl*^+/E397K^ (mean=76.41, *P*=0.0319), *Nefl^E^*^397K/E397K^ (mean=66.49, *P*=0.0001), *Nefl*^+/E397K^ mean compared to *Nefl^E^*^397K/E397K^ mean (*P*=0.0012). (C) Negative area *P*\*=0.0215-0.0445. Wild type (mean=26.55), *Nefl*^+/E397K^ (mean=26.51, NS), *Nefl*^E397K/397K^ (mean=23.93, NS), *Nefl*^+/E397K^ mean compared to *Nefl^E^*^397K/E397K^ mean (NS). A mixed effects analysis with Dunnett’s multiple comparison test and one-way ANOVA with Tukey’s multiple comparison test were used to determine significance. Ms=milliseconds, mV=millivolts, CMAP=compound muscle action potential, NS=not significant, N=number of mice evaluated.

To determine whether there was variability between individual mice in the longitudinal study, consistent with the CMT2E patient population, we analyzed the longitudinal electrophysiology data from P21 to P360 per mouse (**Supplemental Fig. 1A-C**). When measurements were followed for each mouse, there was little variability between P21 wild type mice in distal latency (0.560ms ±0.14); however, there was measurable variability between *Nefl^+/E397K^* (0.870ms ±0.495) and *Nefl^E397K/E397K^* mice (0.844ms ±0.306) (**Fig. 3A**). The variability between *Nefl* mutant mice diminished by P180 (wild type 0.444ms ±0.044; *+/E397K* 0.740ms ±0.119; *E397K/E397K* mice 0.731ms ±0.173); however, distal latency variability was increased in the *Nefl* mutants compared to wild type mice at all timepoints measured (**Supplemental Fig. 1A**). CMAP amplitude and negative area values demonstrated similar variation within cohorts for all genotypes; however, CMAP amplitude values for *Nefl^E397K/E397K^* mice trended lower at all timepoints when compared to wild type mice (**Supplemental Fig. 1B-C**).

## *Nefl* mutant mice showed chronic axonal neuropathy

Our electrophysiology findings showed that there were measurable differences in *Nefl* mutants. To further quantify these deficits, we examined the histopathology of the sciatic nerve at P21, P84, P180, and P360 (**Fig. 4**, **Table 1**). Axon area, axon diameter and G-ratio (ratio of the inner axonal diameter to the total outer diameter) were measured. Cross-sectional images of the sciatic nerve showed significant changes in *Nefl^+/E397K^* and *Nefl^E397K/E397K^* mice at all time points analyzed. At P21, significant changes in axon area were apparent in *Nefl^+/E397K^* (0.0047μm^2^, *P*<0.0001) and *Nefl^E397K/E397K^* (0.0035μm^2^, *P*<0.0001) compared to wild type mice (0.0122μm^2^) **Fig. 4A-E**, **Table 1**). Throughout the analyses, axon area was consistently smaller in *Nefl^+/E397K^* (P360, 0.0148μm^2^, *P*<0.0001) and *Nefl^E397K/E397K^* (P360, 0.0098μm^2^, *P*<0.0001) mice when compared to wild type mice (P360, 0.0314μm^2^) (**Fig. 4A-E**, **Table 1**). Axon diameter was also significantly reduced in *Nefl^+/E397K^* and *Nefl^E397K/E397K^* mice at all time points analyzed (**Fig. 4A-E** **Table 1**). The G-ratio was also reduced in *Nefl^+/E397K^*(P360 0.5557, *P*<0.0001) and *Nefl^E397K/E397K^* mice (P360 0.5515, *P*<0.0001) compared to wild type mice (P360 0.6415). The differences in G-ratio are largely attributed to changes in axon area as we did not observe significant differences in myelination (**Fig. 4A-E**, **Table 1 and not shown**). From three weeks to twelve months, differences in axon area, diameter and G-ratio were more apparent in *Nefl^E397K/E397K^* mice as compared to *Nefl^+/E397K^* mice (**Fig. 4A-E**, **Table 1**). The reduced axon number/field from twelve weeks to six months in *Nefl* mutants largely is not a result of increased number of axons with larger caliber but is attributed to loss of axons (**Fig. 4A-E**, **Table 1**). From six months to twelve months, some regeneration is occurring. There is a substantial increase in the number of axons/field from six months to twelve months in *Nefl^E397K/E397K^* and *Nefl^+/E397K^* mice and those axons represent smaller axons ((**Fig. 4A-E**, **Table 1, Supplemental Fig. 2**). Importantly, the chronic axonal neuropathy observed in *Nefl^E397K/E397K^*and *Nefl^+/E397K^* mice was consistent with the impaired distal latency measured by electrophysiology.

**Figure 4A-D.**
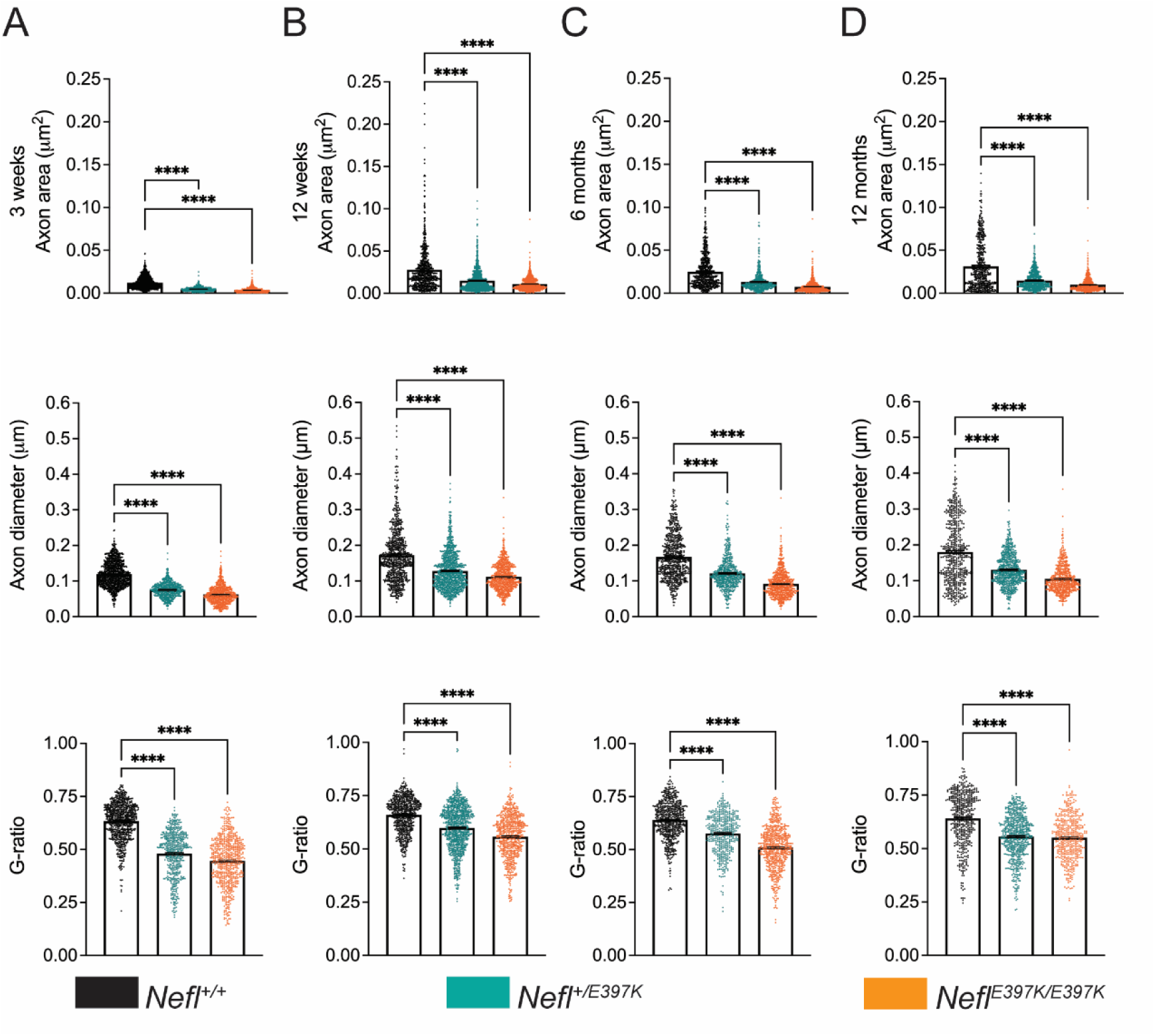
*Nefl* mutant mice showed chronic axonal neuropathy. The sciatic nerve was harvested from wild type (black, N=16) *Nefl*^+/E397K^ (teal, N=17) and *Nefl^E^*^397K/E397K^ (orange, N=16) mice at three weeks, twelve weeks, six months, and twelve months of age. (A) Three weeks axon area for wild type (0.0122μm^2^, 1141 axons), *Nefl*^+/E397K^ (0.0047μm^2^, *P*<0.0001, 623 axons) and *Nefl^E^*^397K/E397K^ (0.0035μm^2^, *P*<0.0001, 724 axons) mice. Three weeks axon diameter for wild type (0.1196μm, 1141 axons), *Nefl*^+/E397K^ (0.0750μm, *P*<0.0001, 623 axons) and *Nefl^E^*^397K/E397K^ (0.0624μm, *P*<0.0001, 724 axons) mice. Three weeks G-ratio for wild type (0.6338, 800 axons), *Nefl*^+/E397K^ (0.4802, *P*<0.0001, 536 axons) and *Nefl^E^*^397K/E397K^ (0.4465, *P*<0.0001, 653 axons) mice. (B) Twelve weeks axon area for wild type (0.0280μm^2^, 759 axons), *Nefl*^+/E397K^ (0.0150μm^2^, *P*<0.0001, 1090 axons) and *Nefl^E^*^397K/E397K^ (0.0110μm^2^, *P*<0.0001, 795 axons) mice. Twelve weeks axon diameter for wild type (0.1724μm, 759 axons), *Nefl*^+/E397K^ (0.1281μm, *P*<0.0001, 1090 axons) and *Nefl^E^*^397K/E397K^ (0.1117μm, *P*<0.0001, 795 axons) mice. Twelve weeks G-ratio for wild type (0.6596, 653 axons), *Nefl*^+/E397K^ (0.5981, *P*<0.0001, 992 axons) and *Nefl^E^*^397K/E397K^ (0.5578, *P*<0.0001, 644 axons) mice. (C) Six months axon area for wild type (0.0250μm^2^, 704 axons), *Nefl*^+/E397K^ (0.0131μm^2^, *P*<0.0001, 553 axons) and *Nefl^E^*^397K/E397K^ (0.0076μm^2^, *P*<0.0001, 769 axons) mice. Six months axon diameter for wild type (0.1670μm, 704 axons), *Nefl*^+/E397K^ (0.1215μm, *P*<0.0001, 553 axons) and *Nefl^E^*^397K/E397K^ (0.0915μm, *P*<0.0001, 769 axons) mice. Six months G-ratio for wild type (0.6386, 622 axons), *Nefl*^+/E397K^ (0.5751, *P*<0.0001, 464 axons) and *Nefl^E^*^397K/E397K^ (0.5093, *P*<0.0001, 635 axons) mice. (D) Twelve months axon area for wild type (0.0314μm^2^, 542 axons), *Nefl*^+/E397K^ (0.0148μm^2^, *P*<0.0001, 719 axons) and *Nefl^E^*^397K/E397K^ (0.0098μm^2^, *P*<0.0001, 627 axons) mice. Twelve months axon diameter for wild type (0.1802μm, 542 axons), *Nefl*^+/E397K^ (0.1302μm, *P*<0.0001, 719 axons) and *Nefl^E^*^397K/E397K^ (0.1049μm, *P*<0.0001, 627 axons) mice. Twelve months G-ratio for wild type (0.6415, 480 axons), *Nefl*^+/E397K^ (0.5557, *P*<0.0001, 636 axons) and *Nefl^E^*^397K/E397K^ (0.5515, *P*<0.0001, 465 axons) mice.Statistics were determined using one-way ANOVA with Dunnett’s multiple comparison test. N=number of mice evaluated.

**Figure 4E.**
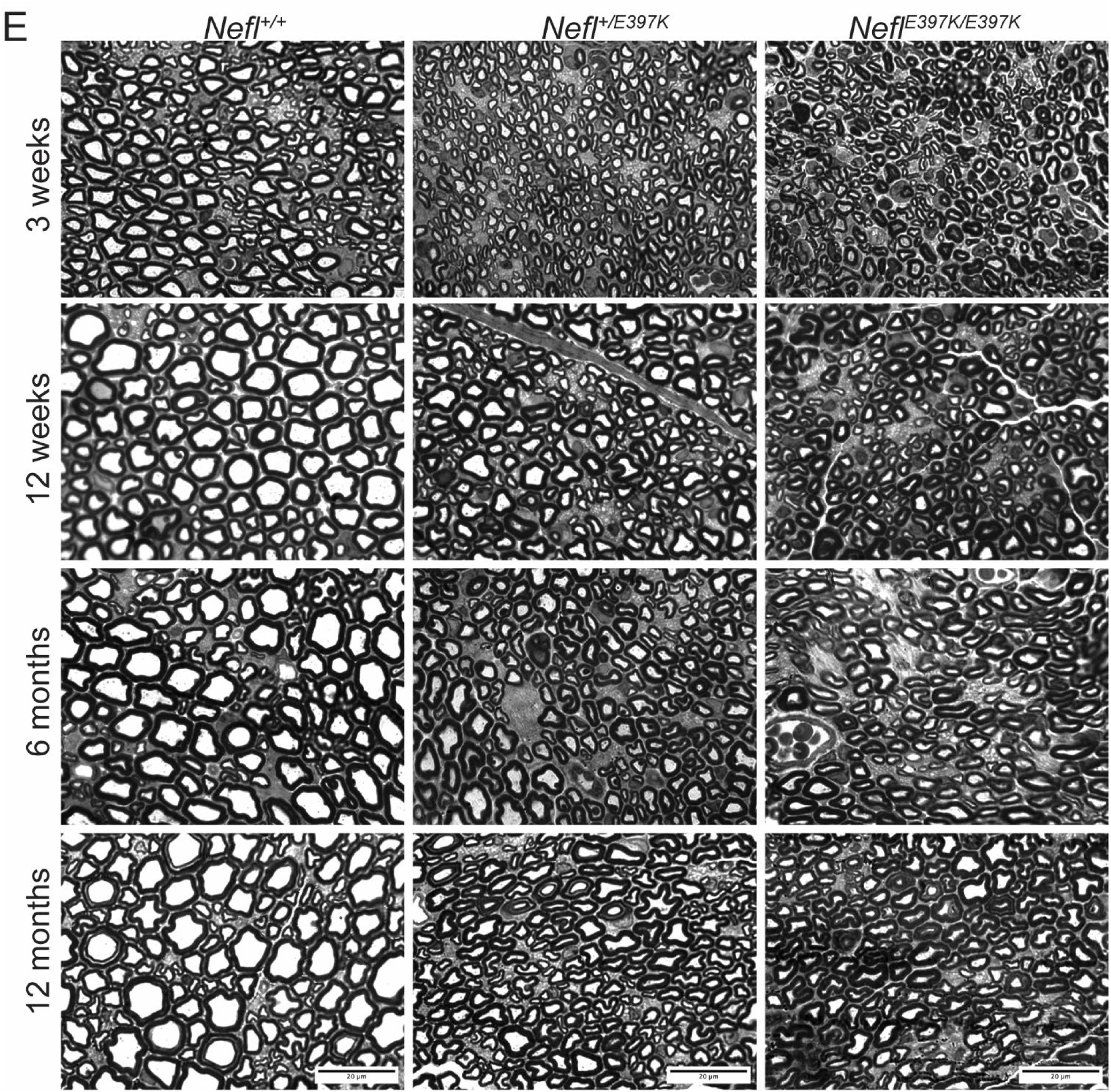
Representative images of axons from wild type, *Nefl*^+/E397K^ and *Nefl^E^*^397K/E397K^ mice at three weeks, twelve weeks, six months and twelve months of age following.

**Table 1.** Axon area, axon diameter, G-ratio and axon numbers.

| Axon area ($\mu\text{m}^2$) | *Nefl*<sup>+/+</sup> | *Nefl*<sup>+/E397K</sup> | *Nefl*<sup>E397K/E397K</sup> |
| --- | --- | --- | --- |
| 3 weeks | 0.0122 | 0.0047 | 0.0035 |
| 12 weeks | 0.0280 | 0.0150 | 0.0110 |
| 6 months | 0.0250 | 0.0131 | 0.0076 |
| 12 months | 0.0314 | 0.0148 | 0.0098 |

| Axon diameter ($\mu\text{m}$) | *Nefl*<sup>+/+</sup> | *Nefl*<sup>+/E397K</sup> | *Nefl*<sup>E397K/E397K</sup> |
| --- | --- | --- | --- |
| 3 weeks | 0.1196 | 0.0750 | 0.0624 |
| 12 weeks | 0.1724 | 0.1281 | 0.1117 |
| 6 months | 0.1670 | 0.1215 | 0.0915 |
| 12 months | 0.1802 | 0.1302 | 0.1049 |

| G-ratio | *Nefl*<sup>+/+</sup> | *Nefl*<sup>+/E397K</sup> | *Nefl*<sup>E397K/E397K</sup> |
| --- | --- | --- | --- |
| 3 weeks | 0.6338 | 0.4802 | 0.4465 |
| 12 weeks | 0.6596 | 0.5981 | 0.5578 |
| 6 months | 0.6386 | 0.5751 | 0.5093 |
| 12 months | 0.6415 | 0.5557 | 0.5515 |

| Average axon number/field | *Nefl*<sup>+/+</sup> | *Nefl*<sup>+/E397K</sup> | *Nefl*<sup>E397K/E397K</sup> |
| --- | --- | --- | --- |
| 3 weeks | 529 | 745 | 731 |
| 12 weeks | 436 | 421 | 548 |
| 6 months | 306 | 354 | 246 |
| 12 months | 285 | 409 | 424 |

When we analyzed axonal area from three weeks to twelve months in *Nefl^+/E397K^* and *Nefl^E397K/E397K^* mice there was a distribution towards more smaller axon calibers with *Nefl^E397K/E397K^* demonstrating more smaller caliber axons than *Nefl^+/E397K^* mice (**Supplemental Fig. 2**). These results are consistent with the electrophysiology studies and axonal pathology.

### Neuromuscular junction denervation was present as disease progressed in *Nefl* mutant mice

To investigate whether neuromuscular junction (NMJ) innervation status was altered in *Nefl* mutant mice, biceps brachii, triceps brachii, gastrocnemius, and tibialis anterior (TA) muscles were analyzed at three weeks, twelve weeks, six months, and twelve months (**Fig. 5** and not shown). At three weeks, there were no significant differences between wild type and *Nefl* mutant mice in any of the muscles examined; however, at twelve weeks the percentage of fully innervated endplates decreased, and partially and fully denervated endplates increased with differences observed between the muscles (**Fig. 5B**). At twelve weeks, the most significant differences were observed in the TA muscle. TA fully innervated endplates in *Nefl^+/E397K^* mice were 88% with 12% partially innervated endplates. *Nefl^E397K/E397K^* mice had 73% fully innervated endplates (*P*=0.0039) with 9% partially innervated and 18% fully denervated endplates in the TA muscle (**Fig. 5B**). The differences in innervation status between wild type and *Nefl* mutants became more apparent at twelve months across all muscles with the greatest differences observed in the TA muscle. TA fully innervated endplates in *Nefl^+/E397K^* mice were 70% (*P*=0.0448) with 24% partially innervated endplates and 6% fully denervated endplates. *Nefl^E397K/E397K^*mice had 56% fully innervated endplates (*P*=0.0013) with 32% partially innervated (*P*=0.0239) and 12% fully denervated endplates in the TA muscle (**Fig. 5C-D**).

**Figure 5A-C:**
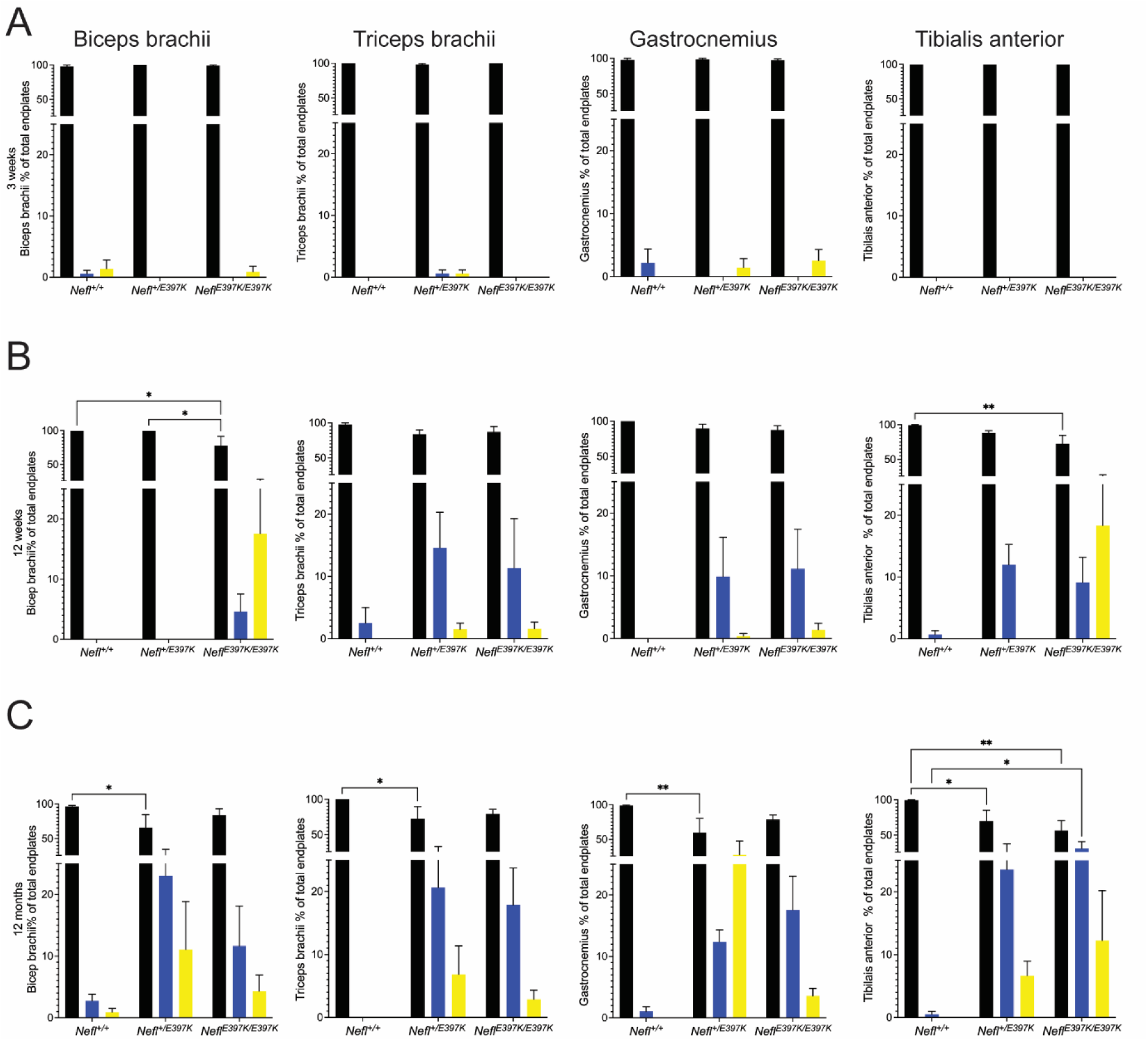
Neuromuscular junction denervation in *Nefl* mutant mice. The neuromuscular junction innervation status of wild type, *Nefl*^+/E397K^, and *Nefl^E^*^397K/E397K^ mice was analyzed at three weeks, twelve weeks, and twelve months of age in the biceps brachii, triceps brachii, gastrocnemius and tibialis anterior (TA) muscles. End plates with complete overlap with the terminal were considered fully innervated (black), end plates with partial overlap were considered partially innervated (blue), and end plates with missing overlapping terminal were considered denervated (yellow). (A) Three-week evaluation. (B) Twelve-week evaluation. Biceps brachii fully innervated endplates between wild type (100%) and *Nefl^E^*^397K/E397K^ mice (78%, *P*=0.0167). Biceps brachii fully innervated endplates between *Nefl*^+/E397K^ and *Nefl^E^*^397K/E397K^ mice (*P*=0.0227). Biceps brachii *Nefl^E397K/397K^* partially innervated endplates 4%, fully denervated endplates 18%. TA fully innervated endplates between wild type (99%) and *Nefl^E^*^397K/E397K^ mice (73%, *P*=0.0039). TA *Nefl^E^*^397K/E397K^ partially innervated endplates 9%, fully denervated endplates 18%. (C) Twelve months evaluation. Biceps brachii fully innervated endplates between wild type (96%) and *Nefl*^+/E397K^ mice (66%, *P*=0.0155). Biceps brachii *Nefl*^+/E397K^ partially innervated endplates 23%, fully denervated endplates 11%. Triceps brachii fully innervated endplates between wild type (100%) and *Nefl*^+/E397K^ mice (73%, *P*=0.0138). Triceps brachii *Nefl*^+/E397K^ partially innervated endplates 21%, fully denervated endplates 6%. Gastrocnemius fully innervated endplates between wild type (99%) and *Nefl*^+/E397K^ mice (60%, *P*=0.0094). Gastrocnemius *Nefl*^+/E397K^ partially innervated endplates 12%, fully denervated endplates 28%. TA fully innervated endplates between wild type (100%) and *Nefl*^+/E397K^ mice (70%, *P*=0.0448). TA *Nefl*^+/E397K^ partially innervated endplates 24%, fully denervated endplates 6%. TA fully innervated endplates between wild type (100%) and *Nefl^E^*^397K/E397K^ mice (56%, *P*=0.0013). TA *Nefl^E^*^397K/E397K^ partially innervated endplates (32%, *P*=0.0239), fully denervated endplates 12%. Four to seven animals were analyzed per muscle. Two-way ANOVA with Tukey’s multiple comparisons test determined statistical significance.

**Figure 5D:**
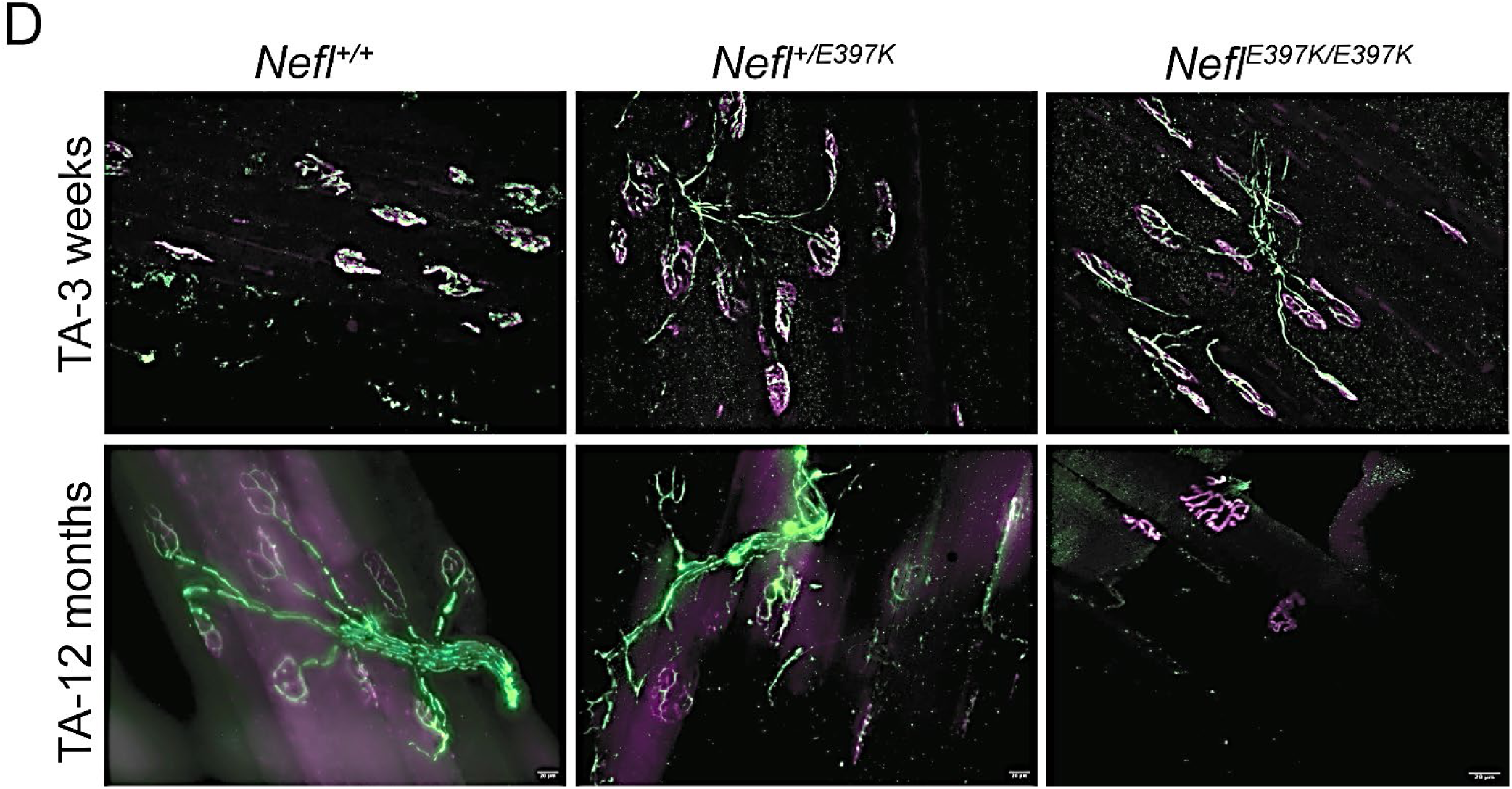
Neuromuscular junction denervation in *Nefl* mutant mice. Representative images for TA muscle at three weeks and twelve months. Anti-neurofilament heavy chain labelled the axon (green) and anti-synaptic vesicle 2 (SV2) (green) labelled the synaptic terminal. Acetylcholine receptors were labeled with Alexa Fluor 594-conjugated α-Bungarotoxin (magenta). Four to seven animals were analyzed per muscle.

Total axon number per field and total NMJ numbers per field were also examined. In wild type mice, the total number of axons per field decreased as the axons grew in area; the total number of NMJs increased (**Fig. 6A-D**, **Table 1**). In P21 *Nefl* mutants, there were more total axons per field than wild type mice as *Nefl* mutant axons were significantly smaller in area; however, total NMJ numbers per field was consistent with wild type mice (**Fig. 6A-D**, **Table 1**). From P21 through P180 there was a consistent decrease in total axons/field in both *Nefl* mutants; however, during this same time there was an increase in total number of NMJ/field consistent with wild type mice (**Fig. 6A-D**, **Table 1**). At P360, *Nefl^E397K/E397K^* axon area was 69% smaller and *Nefl^+/E397K^* axon area was 53% smaller than wild type with 33% and 30% more axons/field, respectively. At P360, the average total number of NMJ/field was 47 for wild type mice, 46 for *Nefl^+/E397K^* mice and 40 for *_Nefl_E397K/E397K* _mice._

**Figure 6.**
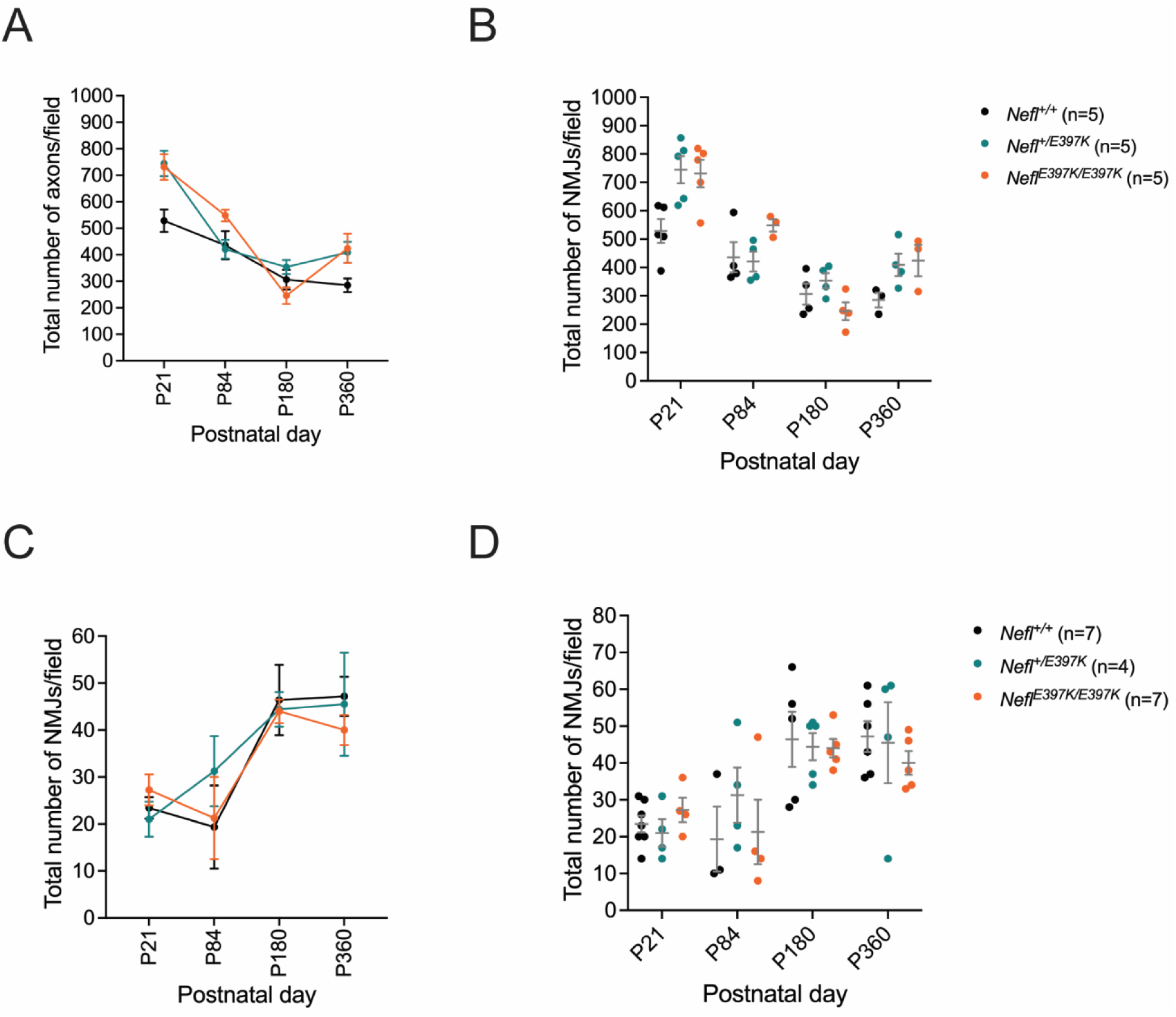
*Nefl* mutants with comparison of axon numbers and NMJs. The total number of axons per 63X field (A, B) and the total neuromuscular junctions (NMJs) per 20X field (C, D) were quantified and compared. Wild type (black), *Nefl*^+/E397K^ (teal), and *Nefl^E^*^397K/E397K^ (orange) mice were analyzed at three weeks, twelve weeks, six months, and twelve months of age. Axons of the sciatic nerve and NMJs associated with the gastrocnemius muscle were quantified. Wild type (P360=285 axons, 47=NMJ), *Nefl*^+/E397K^ (P360=409 axons, 46=NMJ), and *Nefl^E^*^397K/E397K^ (P360=424 axons, 40=NMJ) mice. P=postnatal day, n=number of animals evaluated.

## Discussion

Here we report two *Nefl-E397K* mouse models that present with early electrophysiology differences and axon pathology that persisted throughout the lifespan of the mice. Additionally, in the characterization of *Nefl^+/E397K^*or *Nefl^E397K/E397K^* mice we found that *Nefl^E397K/E397K^*mice demonstrate disease pathology earlier than *Nefl^+/E397K^* mice and in most instances disease pathology was more severe. Consistent with CMT2E patients, there was not a reduction in lifespan for *Nefl^+/E397K^* or *Nefl^E397K/E397K^*mice; however, weight for *Nefl^+/E397K^*mice trended upward while weight for *Nefl^E397K/E397K^*mice trended downward. These differences might be reflected by the mobility of the mice, any associated neuropathic pain and disease severity.

Electrophysiology measurements recorded with individual cohorts or within the longitudinal study were consistent demonstrated significant differences as early as P21, revealing an early, quantifiable phenotype. Distal latency was significantly prolonged throughout the 360 days while P-P CMAP amplitude was consistently lower in the *Nefl* mutants. At three weeks, electrophysiology measurements (distal latency, P-P CMAP amplitude and negative area) demonstrated significant differences from wild type cohorts. A possible interpretation of these results would be that there was a decrease in the axon caliber (increased distal latency), reduced number of motor units activated (CMAP) and reduced activated muscle fiber (negative area). The distal latency was consistent with the significant decrease in axon area and diameter observed in *Nefl^+/E397K^* and *Nefl^E397K/E397K^* mice. At three weeks, there was no NMJ denervation present in the forelimb nor hindlimb muscles examined in *Nefl* mutants; however, there were more and smaller axons with fewer total NMJs consistent with the reduction in activated motor units. In our companion study, significantly reduced muscle fiber area in *Nefl* mutants further supports the electrophysiology findings.

Interestingly, from three to twelve weeks distal latency was further prolonged in *Nefl* mutant mice with *Nefl*^E397K/E397K^ more severe than *Nefl*^+/E397K^ mice. There was some improvement in P-P CMAP amplitude and negative area at twelve weeks; however, both remained statistically different from the wild type cohort. *Nefl* mutant axonal growth continued, but axon area remained significantly smaller than wild type axons contributing to the prolonged distal latency. For *Nefl*^E397K/E397K^ mice from three weeks to twelve weeks, there was a three-fold increase in axon area; however, axons were 39% the size of wild type axons. *Nefl^+/E397K^* mice presented with more, larger axons than *Nefl^E397K/E397K^* mice with increased number of total NMJs. Overall, NMJ innervation was largely preserved in *Nefl^+/E397K^* mice with increased partial and full denervated endplates observed in *Nefl^E397K/E397K^* mice. NMJ number/field were similar between wild type and *Nefl^E397K/E397K^* mice; however, *Nefl^E397K/E397K^* mice had more and smaller axons. The increased axon caliber, “maintenance” of NMJ innervation and potential for sprouting likely stabilized P-P CMAP and negative area in the *Nefl* mutants.

From twelve weeks to six months, axon degeneration persisted in the *Nefl* mutants, most notably in *Nefl^E397K/E397K^*mice. *Nefl^+/E397K^* demonstrated a 13% reduction in axon caliber with a 16% reduction in the number of axons per field (smaller axons and fewer axons counted). *Nefl^E397K/E397K^* mice demonstrated a 31% reduction in axon caliber with a 55% reduction in the number of axons (much smaller axons and significantly fewer axons counted). Additionally, there was an increase in the absence of axons within the fields scored at six months indicating axon loss. *Nefl^E397K/E397K^* mice had 70% smaller axon area than wild type mice at six months and a 20% reduction in the number of axons. Notably, *Nefl* mutant mice increased total NMJs consistent with wild type mice suggesting compensatory mechanisms such as sprouting were occurring. Together, these results support the electrophysiology; distal latency was prolonged due to the small axons. P-P CMAP and negative area were largely preserved due to compensatory mechanisms that maintained NMJ innervation and facilitated sprouting as well as muscle fiber hypertrophy in *Nefl* mutant mice (companion manuscript).

From six months to twelve months, axon caliber slightly increased in *Nefl* mutant mice; however, wild type mice demonstrated a decrease in the number of axons/field (consistent with increased caliber) while *Nefl* mutants demonstrated increased axon number (*Nefl^+/E397K^*13%, *Nefl^E397K/E397K^* 42%) suggesting regeneration was occurring. More, smaller axons were present at twelve months in comparison to six months in *Nefl* mutants. The increased but smaller axons resulted in significantly prolonged distal latency. While there was not a significant difference in P-P CMAP amplitude and negative area, *Nefl* mutants consistently had lower measurements than their wild type cohort. The maintenance of NMJ numbers could serve as a compensatory mechanism for the changes in NMJ innervation status.

## Materials and Methods

### Animals

All experimental procedures were approved by the University of Missouri Animal Care and Use committee and were performed according to the guidelines set forth in the Guide for the Use and Care of Laboratory Animals.

### CRISPR/Cas9, *Nefl*-*E397K* sgRNA and repair template

*Nefl* mice were generated on a C57BL/6 background using CRISPR technology at the University of Missouri Animal Modeling Core. An enhanced-specificity Cas9 (eSPCas9) protein was used to reduce off-target effects. Any predicted off-target site with less than a 2bp mismatch (including DNA or RNA bulges) or with less than 3bp mismatches if no mismatches are in the 12bp seed region of the sgRNA was PCR amplified and sequenced to ensure no erroneous edits were made. The sgRNA was ordered as chemically modified synthetic sgRNA (Synthego). The repair template was chemically synthesized as a 199bp single stranded DNA oligo (ssODN) (IDT). The ssODN was complementary to the non-target strand and contained symmetrical homology arms. sgRNA sequence: 5’-TCTTGGAAGGCGAAGAGACC-3’. Repair template sequence (sgRNA sequence disrupted by the desired mutation) 5’-GAGGAAAGTAATGAATGTGGGCTTAGAGCAATGAACACATCCAGCCTTGCTCTAACTGTAC TCTTCATTCCCTCTCCACCAGAAAACTCTTGGAAGGC**A**AAGAGACCAGACTCAGTTTCACC

AGCGTGGGTAGCATAACCAGCGGCTACTCTCAGAGCTCGCAGGTCTTCGGCCGTTCTGCTTACAGTGGCTTGCAGAGC −3’. The boxed area represents the sgRNA target sequence, and the bold underlined letter indicates the mutation engineered in the DNA repair template.

### Zygote electroporation and embryo transfer

A mix containing a final concentration of 3.0μM sgRNA, 2.0μM Cas9 protein (Sigma) and 1.6μM ssODN was made immediately prior to electroporation. CRISPR sgRNA/Cas9 RNP complexes were formed with incubation at room temperature for 10 minutes followed by the addition of the ssODN repair template. Zygotes were electroporated (NepaGene21) using a 1.5mm gap glass slide electrode under the following conditions: Poring pulse: 40V, 3.5ms length, 50ms interval, 10% decay rate, positive polarity (x4 pulses) Transfer pulse: 5V, 50ms length, 50ms interval, 40% decay rate, alternating polarity (x5 pulses). Electroporated zygotes were transferred to pseudo pregnant surrogate females.

### Genotyping

Genotyping of neonatal pups was performed at postnatal day (P) 0. Genomic DNA isolation was performed using a protocol from Jacksons labs. C57BL/6-*Nefl^E397K^* mice were genotyped using a high-resolution melt analysis (EvaGreen qPCR precision melt supermix, Bio-Rad) and the primers 5’-CTTCATTCCCTCTCCACCAG-3’ and 5’-CACGCTGGTGAAACTGAG-3’. PCR conditions were 95°C denaturing for 2:00 minutes (m) followed by 39 cycles of 95°C denaturing for 10 seconds (s), 60°C annealing for 30s and 72°C extension for 30s, 95°C for 30s, 60°C for 1:00m, and melt curve 65°C to 95°C in increments of 0.2 every 10s. Melting temperature for the wild type, *Nefl^+/E397K^*, and *Nefl^E397K/E397K^* were 78.5 ± 0.03°C, 77.7 ± 0.03°C, and 77.2 ± 0.06°C, respectively.

### Electrophysiology studies

Measurements of the right gastrocnemius muscle were recorded following stimulation of the sciatic nerve as previously described (25). Briefly, mice were anesthetized with isoflurane (2-5% for induction and 2-3% for maintenance) and placed on a warming mat set at 37°C. The right hindlimb was shaved and an active ring electrode was placed over the gastrocnemius muscle and a reference ring electrode was placed over the metatarsals of the right hind paw (Alpine Biomed, Skovlunde, Denmark). Spectra 360 electrode gel was applied to decrease impedance (Parker Laboratories, Fairfield, NJ). A common reference electrode was placed around the tail. Two 28-gauge monopolar needle electrodes (Teca, Oxford Instruments Medical, New York, NY) were placed on each side of the sciatic nerve in the region of the proximal thigh. A portable electrodiagnostic system (Cadwell Sierra Summit, Kennewick, WA) was used to stimulate the sciatic nerve (0.1ms pulse, 1–10mA intensity) and distal latency, compound muscle action potential amplitude, and negative area were recorded.

### Sciatic nerve dissection and processing

Sciatic nerves were fixed with 8% glutaraldehyde in phosphate buffer and processed for resin (Poly/Bed® 812; #21844-1; Polysciences Inc.) embedding as previously described (17, 26). After fixation, nerves were incubated in 2% osmium tetroxide in phosphate buffer followed by rinses in ascending ethanol concentrations (50%, 70%, 80%, 95%, and 100%) and propylene oxide. Samples were then incubated in a 1:1 propylene oxide: resin mixture for 1 hour, incubated in resin overnight, placed in a resin mold, and cured at 60°C for 8 hours. Semi-thin sections of 1μm were stained with alkaline toluidine blue, cover-slipped with Permount mounting medium (ThermoFisher Scientific), and visualized by light microscopy (Leica DM5500 B, Leica Microsystems Inc.). Image quantification was performed in a blind manner using the semi-automated MyelTracer software (27). Axon area, perimeter, diameter, and G-ratios were recorded.

### Neuromuscular junction (NMJ) immunohistochemistry

Mice were perfused with 4% PFA. NMJ immunohistochemistry whole mount preparations were stained and analyzed as previously described (28). Muscles were incubated with primary antibodies anti-Neurofilament Heavy Chain (NF-H) (1:2000; AB5539, Chemicon, EMD Millipore) and anti-Synaptic Vesicle 2 (SV2) (1:200; YE269, Life Technologies) followed by Donkey anti-Chicken Alexa Fluor 488 (1:400; Jackson ImmunoResearch) and Goat anti-Rabbit Alexa Fluor 488 (1:200; Jackson ImmunoResearch) secondary antibodies to label the axon and synaptic terminal. Acetylcholine receptors were labeled with Alexa Fluor 594-conjugated α-Bungarotoxin (1:200; Life Technologies). NMJ analyses were performed blind on three randomly selected fields at 20X magnification (Leica DM5500 B, Leica Microsystems Inc.). Analyses were based on the overlap of the end plate and synaptic terminal. End plates with complete overlap with the terminal were considered fully innervated, end plates with partial overlap were considered partially innervated, and end plates with missing overlapping terminal were considered denervated using Fiji software (NIH). Representative images were obtained using a laser scanning confocal microscope at 40x magnification (Leica TCS SP8, Leica Microsystems Inc.).

### Statistical analyses

Statistical analyses for each experimental protocol are noted within the figure legends.

## Acknowledgements

We acknowledge the MU Animal Modeling Core, MU Genomics Technology Core, and MU Advanced Light Microscopy Core for assistance with these studies.

## Author Contributions

DPL: designed research studies, conducted experiments, acquired data, analyzed data, prepared manuscript, edited manuscript

AS: conducted experiments, acquired data, analyzed data

FLT: conducted experiments, acquired data, analyzed data

MA: conducted experiments, acquired data, analyzed data

WA: acquired data, analyzed data

MAL: conducted experiments, acquired data, analyzed data, edited manuscript

CLL: designed research studies, analyzed data, edited manuscript

---

**Supplemental Figure 1.**
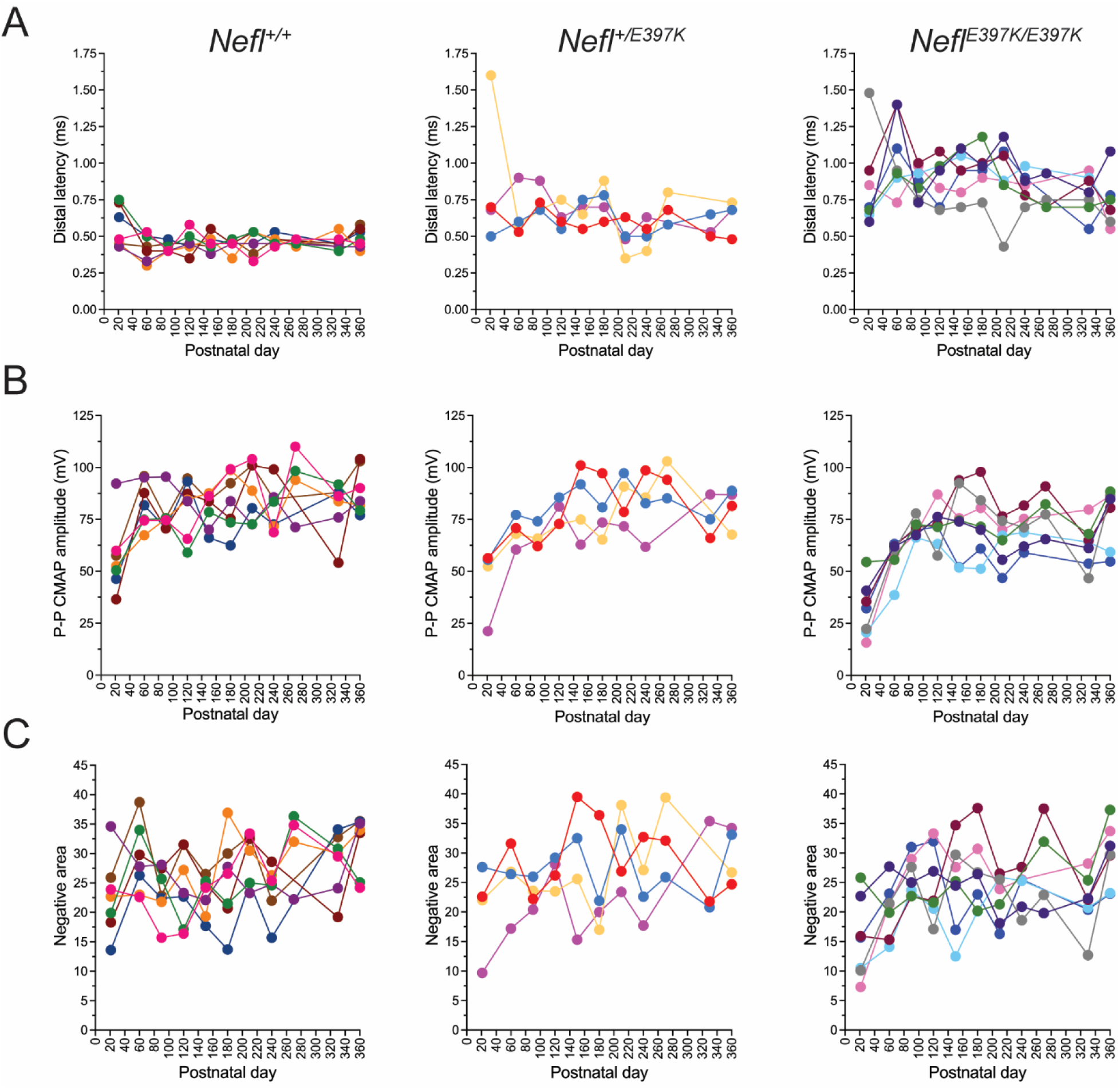
Longitudinal study evaluated per mouse. Distal latency, Peak to Peak (P-P) CMAP amplitude and negative area were measured following stimulation of the sciatic nerve and recordings from the gastrocnemius muscle. Wild type (N=7), *Nefl*^+/E397K^ (N=4) and *Nefl^E^*^397K/E397K^ (N=7) mice were evaluated at P21, P60, P90, P120, P150, P180, P210, P240, P270, P330 and P360 days. Each color represents one mouse (A) Distal latency. (B) P-P CMAP amplitude. (C) Negative area. Ms=milliseconds, mV=millivolts, CMAP=compound muscle action potential, N=number of mice evaluated.

**Supplemental Figure 2.**
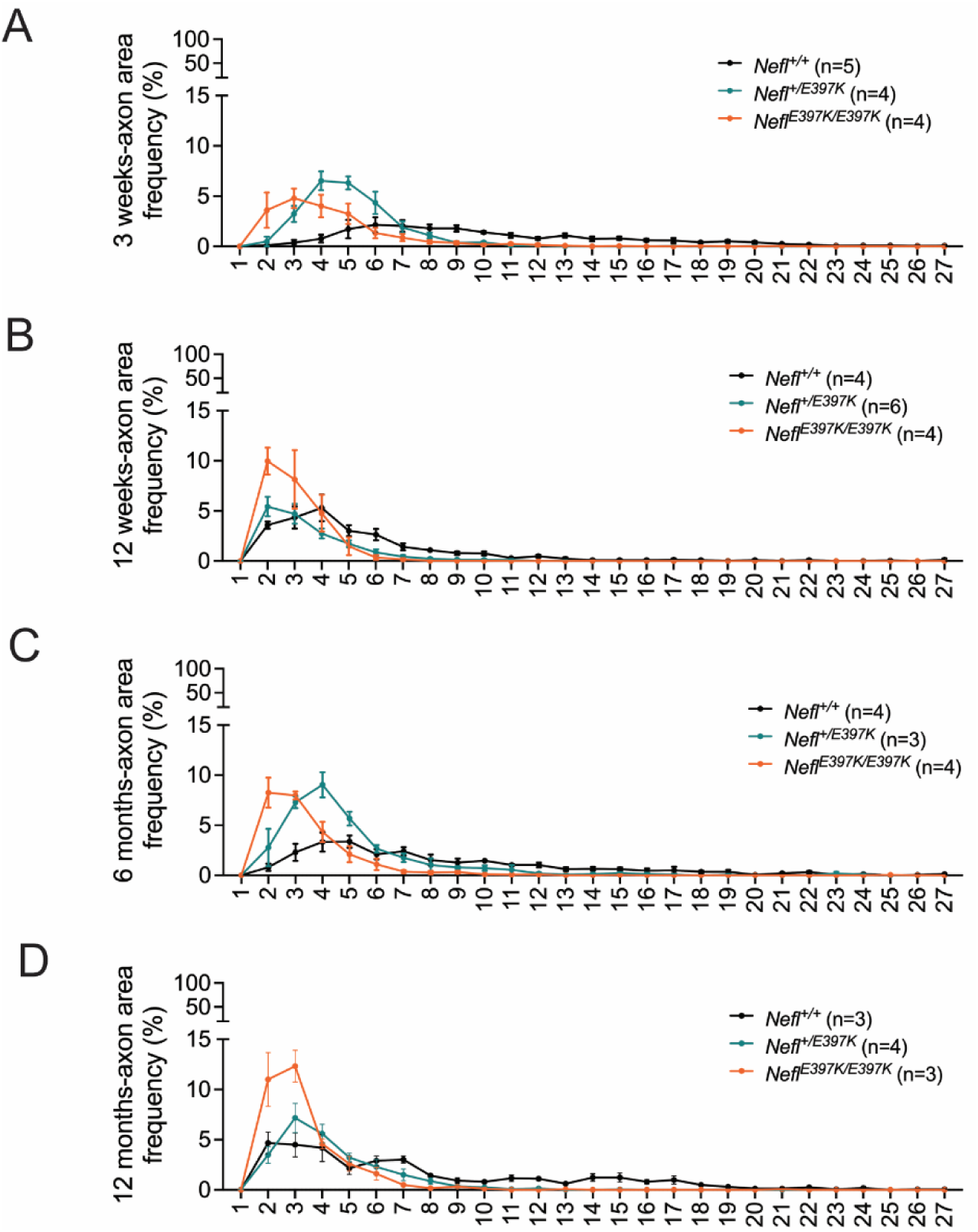
Axon area distribution per age group. The sciatic nerve was harvested from wild type, *Nefl*^+/E397K^, and *Nefl^E^*^397K/E397K^ mice at three weeks, twelve weeks, six months, and twelve months of age. N=number of mice evaluated. Three-week values in μm^2^ 1=0-1.4, 2=1.4-15.06, 3=15.06-28.71, 4=28.71-42.37, 5=42.37-56.02, 6=56.02-69.67, 7=69.97-83.33, 8=83.33-96.98, 9=96.98-110.64, 10=110.64-124.29, 11=124.29-137.94, 12=137.94-151.60, 13=151.60-165.25, 14=165.25-178.91, 15=178.91-192.56, 16=192.56-206.21, 17=206.21-219.87, 18=219.87-233.52, 19=233.52-247.17, 20=247.17-260.83, 21=260.83-274.48, 22=274.48-288.14, 23=288.14-301.79, 24=301.79-315.44, 25=315.44-329.10, 26=329.10-342.75, 27=>342.75. Twelve-week values in μm^2^ 1=0-6.87, 2=6.87-74.59, 3=74.59-142.32, 4=142.32-210.04, 5=210.04-277.77, 6=277.77-345.49, 7=345.59-413.22, 8=413.22-480.94, 9=480.94-548.67, 10=548.67-616.39, 11=616.39-648.12, 12=648.12-751.84, 13=751.84-819.57, 14=819.57-887.29, 15=887.29-955.05, 16=955.05-1022.74, 17=1022.74-1090.47, 18=1090.47-1158.19, 19=1158.19-1225.92, 20=1225.92-1293.64, 21=1293.64-1361.37, 22=1361.37-1429.09, 23=1429.09-1496.81, 24=1496.81-1564.54, 25=1564.54-1632.26, 26=1632.26-1699.99, 27=>1699.99. Six-month values in μm^2^ 1=0-3.59, 2=3.59-40.36, 3=40.36-77.13, 4=77.13-113.89, 5=113.89-150.66, 6=150.66-187.43, 7=187.43-224.20, 8=224.20-260.96, 9=260.96-297.73, 10=297.73-334.50, 11=334.50-371.27, 12=371.27-408.03, 13=408.03-444.80, 14=440.80-481.57, 15=481.57-518.34, 16=518.34-555.10, 17=555.10-591.87, 18=591.87-628.64, 19=628.64-665.41, 20=665.41-702.17, 21=702.17-738.94, 22=738.94-775.51, 23=775.51-812.48, 24=812.48-849.24, 25=849.24-886.01, 26=886.01-922.78, 27=>922.78. Twelve-month values in μm^2^ 1=0-3.59, 2=3.59-55.54, 3=55.54-107.50, 4=107.50-159.45, 5=159.45-211.40, 6=211.40-263.35, 7=263.35-315.31, 8=315.31-367.26, 9=367.26-419.21, 10=419.21-471.17, 11=471.17-523.12, 12=523.12-575.07, 13=575.07-627.02, 14=627.02-678.98, 15=678.98-730.93, 16=730.93-782.88, 17=782.88-834.84, 18=834.84-886.79, 19=886.79-938.74, 20=938.74-990.69, 21=990.69-1042.64, 22=1042.64-1094.50, 23=1094.50-1146.55, 24=1146.55-1198.51, 25=1198.51-1250.46, 26=1250.46-1302.41, 27=>1302.41.

## References

1. Mersiyanova, I.V., Perepelov, A.V., Polyakov, A.V., Sitnikov, V.F., Dadali, E.L., Oparin, R.B., Petrin, A.N. and Evgrafov, O.V. (2000) A new variant of Charcot-Marie-Tooth disease type 2 is probably the result of a mutation in the neurofilament-light gene. Am. J. Hum. Genet., 67, 37–46.

2. 2. Della Marina, A., Hentschel, A., Czech, A., Schara-Schmidt, U., Preusse, C., Laner, A., Abicht, A., Ruck, T., Weis, J., Choueiri, C., et al. (2024) Novel Genetic and Biochemical Insights into the Spectrum of NEFL-Associated Phenotypes. J. Neuromuscul. Dis., 11, 625–645.

3. Stone, E.J., Kolb, S.J. and Brown, A. (2021) A review and analysis of the clinical literature on Charcot-Marie-Tooth disease caused by mutations in neurofilament protein L. Cytoskeleton (Hoboken), 78, 97–110.

4. 4. De Jonghe, P., Mersivanova, I., Nelis, E., Del Favero, J., Martin, J.J., Van Broeckhoven, C., Evgrafov, O. and Timmerman, V. (2001) Further evidence that neurofilament light chain gene mutations can cause Charcot-Marie-Tooth disease type 2E. Ann. Neurol., 49, 245–249.

5. 5. Jordanova, A., De Jonghe, P., Boerkoel, C.F., Takashima, H., De Vriendt, E., Ceuterick, C., Martin, J.J., Butler, I.J., Mancias, P., Papasozomenos, S., et al. (2003) Mutations in the neurofilament light chain gene (NEFL) cause early onset severe Charcot-Marie-Tooth disease. Brain, 126, 590–597.

6. Elbracht, M., Senderek, J., Schara, U., Nolte, K., Klopstock, T., Roos, A., Reimann, J., Zerres, K., Weis, J. and Rudnik-Schoneborn, S. (2014) Clinical and morphological variability of the E396K mutation in the neurofilament light chain gene in patients with Charcot-Marie-Tooth disease type 2E. Clin. Neuropathol., 33, 335–343.

7. Herrmann, H. and Aebi, U. (2016) Intermediate Filaments: Structure and Assembly. Cold Spring Harb. Perspect. Biol., 8.

8. Herrmann, H., Strelkov, S.V., Burkhard, P. and Aebi, U. (2009) Intermediate filaments: primary determinants of cell architecture and plasticity. J. Clin. Invest., 119, 1772–1783.

9. Eriksson, J.E., Dechat, T., Grin, B., Helfand, B., Mendez, M., Pallari, H.M. and Goldman, R.D. (2009) Introducing intermediate filaments: from discovery to disease. J. Clin. Invest., 119, 1763–1771.

10. Liem, R.K. and Messing, A. (2009) Dysfunctions of neuronal and glial intermediate filaments in disease. J. Clin. Invest., 119, 1814–1824.

11. Kotaich, F., Caillol, D. and Bomont, P. (2023) Neurofilaments in health and Charcot-Marie-Tooth disease. Front. Cell Dev. Biol., 11, 1275155.

12. Pisciotta, C., Bai, Y., Brennan, K.M., Wu, X., Grider, T., Feely, S., Wang, S., Moore, S., Siskind, C., Gonzalez, M. et al. (2015) Reduced neurofilament expression in cutaneous nerve fibers of patients with CMT2E. Neurology, 85, 228–234.

13. Brownlees, J., Ackerley, S., Grierson, A.J., Jacobsen, N.J., Shea, K., Anderton, B.H., Leigh, P.N., Shaw, C.E. and Miller, C.C. (2002) Charcot-Marie-Tooth disease neurofilament mutations disrupt neurofilament assembly and axonal transport. Hum. Mol. Genet., 11, 2837–2844.

14. Lee, M.K., Marszalek, J.R. and Cleveland, D.W. (1994) A mutant neurofilament subunit causes massive, selective motor neuron death: implications for the pathogenesis of human motor neuron disease. Neuron, 13, 975–988.

15. Perez-Olle, R., Leung, C.L. and Liem, R.K. (2002) Effects of Charcot-Marie-Tooth-linked mutations of the neurofilament light subunit on intermediate filament formation. J. Cell Sci., 115, 4937–4946.

16. 16. Adebola, A.A., Di Castri, T., He, C.Z., Salvatierra, L.A., Zhao, J., Brown, K., Lin, C.S., Worman, H.J. and Liem, R.K. (2015) Neurofilament light polypeptide gene N98S mutation in mice leads to neurofilament network abnormalities and a Charcot-Marie-Tooth Type 2E phenotype. Hum. Mol. Genet., 24, 2163–2174.

17. Lancaster, E., Li, J., Hanania, T., Liem, R., Scheideler, M.A. and Scherer, S.S. (2018) Myelinated axons fail to develop properly in a genetically authentic mouse model of Charcot-Marie-Tooth disease type 2E. Exp. Neurol., 308, 13–25.

18. Shen, H., Barry, D.M., Dale, J.M., Garcia, V.B., Calcutt, N.A. and Garcia, M.L. (2011) Muscle pathology without severe nerve pathology in a new mouse model of Charcot-Marie-Tooth disease type 2E. Hum. Mol. Genet., 20, 2535–2548.

19. Dale, J.M., Villalon, E., Shannon, S.G., Barry, D.M., Markey, R.M., Garcia, V.B. and Garcia, M.L. (2012) Expressing hNF-LE397K results in abnormal gaiting in a transgenic model of CMT2E. Genes Brain Behav., 11, 360–365.

20. Dequen, F., Filali, M., Lariviere, R.C., Perrot, R., Hisanaga, S. and Julien, J.P. (2010) Reversal of neuropathy phenotypes in conditional mouse model of Charcot-Marie-Tooth disease type 2E. Hum. Mol. Genet., 19, 2616–2629.

21. Filali, M., Dequen, F., Lalonde, R. and Julien, J.P. (2011) Sensorimotor and cognitive function of a NEFL(P22S) mutant model of Charcot-Marie-Tooth disease type 2E. Behav. Brain Res., 219, 175–180.

22. Zhao, J., Brown, K. and Liem, R.K.H. (2017) Abnormal neurofilament inclusions and segregations in dorsal root ganglia of a Charcot-Marie-Tooth type 2E mouse model. PLoS One, 12, e0180038.

23. Miltenberger-Miltenyi, G., Janecke, A.R., Wanschitz, J.V., Timmerman, V., Windpassinger, C., Auer-Grumbach, M. and Loscher, W.N. (2007) Clinical and electrophysiological features in Charcot-Marie-Tooth disease with mutations in the NEFL gene. Arch. Neurol., 64, 966–970.

24. 24. Bienfait, H.M., Verhamme, C., van Schaik, I.N., Koelman, J.H., de Visser, B.W., de Haan, R.J., Baas, F., van Engelen, B.G. and de Visser, M. (2006) Comparison of CMT1A and CMT2: similarities and differences. J. Neurol., 253, 1572–1580.

25. Arnold, W.D., Sheth, K.A., Wier, C.G., Kissel, J.T., Burghes, A.H. and Kolb, S.J. (2015) Electrophysiological Motor Unit Number Estimation (MUNE) Measuring Compound Muscle Action Potential (CMAP) in Mouse Hindlimb Muscles. J. Vis. Exp.

26. Rich, K.A., Pino, M.G., Yalvac, M.E., Fox, A., Harris, H., Balch, M.H.H., Arnold, W.D. and Kolb, S.J. (2023) Impaired motor unit recovery and maintenance in a knock-in mouse model of ALS-associated Kif5a variant. Neurobiol. Dis., 182, 106148.

27. Kaiser, T., Allen, H.M., Kwon, O., Barak, B., Wang, J., He, Z., Jiang, M. and Feng, G. (2021) MyelTracer: A Semi-Automated Software for Myelin g-Ratio Quantification. eNeuro., 8.

28. 28. Smith, C.E., Lorson, M.A., Ricardez Hernandez, S.M., Al Rawi, Z., Mao, J., Marquez, J., Villalon, E., Keilholz, A.N., Smith, C.L., Garro-Kacher, M.O., et al. (2022) The Ighmbp2D564N mouse model is the first SMARD1 model to demonstrate respiratory defects. Hum. Mol. Genet., 31, 1293–1307.

